# Evidence for a central ecological function of predatory Myxococcota in soil ecosystems

**DOI:** 10.64898/2026.10.21.760627

**Authors:** Khalid El Karkouri, Benoit Perez-Lamarque, Tâm Mignot

## Abstract

Myxococcota is a recently classified phylum including bacteria renowned for their predatory lifestyle. However, ecological functions of these bacteria remain under studied. Here, we examined the ecological position and potential impact of Myxococcota communities across diverse soil habitats using the global geographical collection of 16S microbial communities structured by the Earth Microbiome Project (EMP). First, we found that the abundance of Myxococcota varied greatly from 6.3% to 0.8% across 13 soil habitats. A deeper microbial analysis in two selected habitats showed that Myxococcota was highly abundant (4.9%) ranking 5 in field soil (a grassland planted with switchgrass) than in sand biofilter (an engineered system for water quality production, 1.3% and 11, resp.). Moreover, both Myxococcota and microbial communities displayed a significantly higher α-diversity in field soil than in sand biofilter environments, suggesting that diversity of Myxococcota seems to be associated with that of microbial communities in these habitats. Using co-occurrence network analysis, we found that Myxococcota exhibited a high eigenvector centrality (5.1%) and one hub taxon, Polyangiales, in field soil habitat. In contrast, Myxococcota centrality was less pronounced (1.1%) in the sand biofilter, where consistent negative correlations were also detected. Altogether, these results suggest that Myxococcota communities experienced beneficial interactions in field soil environment, whereas they may have faced competitive relationships in sand biofilter habitat. Overall, our findings provide further insights into the status of Myxococcota communities in soil environments, suggesting their fundamental roles in the structure of soil microbial communities.

**IMPORTANCE:** Myxococcota is a bacterial phylum characterized by a predatory lifestyle in soil microbial ecosystems and exhibits a high genomic and metabolic diversity. However, the ecology of Myxococcota is understudied and remains an important challenge since several studies on predation mechanisms, regulation of microbial communities, or protection against soil-borne phytopathogens were mainly investigated on a subset of Myxococcota taxa under laboratory or greenhouse conditions. The current study showed that Myxococcota is taxonomically diverse in soil habitats. It can interact with distinct patterns of bacterial taxa and may be one of the most central drivers or hubs among microbial communities in soil environments. These findings would advance our understanding of Myxococcota diversity and ecology as well as their potential impact on regulating microbial communities in soil ecosystems which are important for developing crop protection strategies.

## INTRODUCTION

Soil ecosystems are associated with complex microbial communities that are growing increasingly recognized as the dominant drivers of soil ecology, from plant fitness to decomposition, nutrient-cycling etc. (1, 2). The soil microbial food web is regulated by microbial predator-prey interactions, the complexity of which is only emerging (3). Recent explorations of the prevalence of active microbial soil predators, bacteria but also protists have revealed that perhaps the most represented soil predators are bacteria of the Myxococcota phylum (3). These bacteria are widely distributed across diverse environments such as soil, sand, plant, rhizosphere and water (4–6). Furthermore, RNA-Sip analyses converged in showing that they are the most active predators in diverse soil but also sludges from water treatment plants. Their ubiquitous distribution suggests an important ecological role, potentially positioning members of the Myxococcota as keystone species whose presence could critically influence nutrient cycling and the diversity and stability of the ecosystem (3).

Myxococcota is a phylum including bacteria renowned for their predatory lifestyle. Formerly ranged within the class Deltaproteobacteria, it was reclassified and validated by the International Committee on Systematics of Prokaryotes (ICSP) as a distinct phylum based on phylogenetic evidence (7,8). Despite growing interest, morphological characterization and taxonomic identification of Myxococcota individuals remain challenging, and new genera and species within this lineage continue to be discovered (5, 9, 10). This difficulty is largely due to their substantial genomic diversity. Members of the phylum exhibit highly variable genome sizes—ranging from 4.3 to 16.04 Mbp—and GC contents spanning 52% to 75% (https://www.ncbi.nlm.nih.gov/datasets/genome). To date, only a subset of the Myxococcota—primarily *Myxococcus xanthus* of the Myxococcaceae family—has been extensively studied under laboratory conditions. Decades of research have shown that these bacteria hunt cooperatively, swarming into their prey and killing them through direct contact and the secretion of secondary metabolites to digest them extracellularly (11–18). Remarkably, under nutrient-deprived conditions, they form multicellular fruiting bodies that contain spores capable of remaining dormant until favorable conditions or prey availability returns (12).

Although pure culture studies in laboratory context have provided invaluable information on predatory mechanisms, they do not provide information on the interactions that take place in the actual ecological niches. In this respect, perhaps the closest experimental demonstration that myxobacterial predation contributes to the regulation of microbial communities was its ability to promote protection against soil-borne phytopathogens. For example, *M. xanthus* effectively suppressed Tomato Bacterial Wilt (TBW) by preying on *Ralstonia solanacearum* in greenhouse experiments (19). Similarly, *Corallococcus*, another myxobacterium, reduced *Fusarium* infection in cucumbers (20). In that case, the presence of *Corallococcus* was associated with profound changes in the cucumber-associated microbial community, suggesting that bacterial predation plays a significant role in shaping this ecosystem (20). These investigations were further supported by a study in which the addition of *Myxococcus* to a 16-species soil microcosm impacted the microbial community both positively and negatively (21).

Demonstrating the ecological importance of Myxococcota is therefore an important challenge since *in vitro* studies do not also include the complexity of soil microbial communities. A major problem also being that soil communities are largely composed of unculturable microbes which is mainly due to a lack of knowledge of their biology and ecology as well as to the use of inappropriate culture conditions (22, 23). In the metagenomic era, public repositories databases and the Earth Microbiome Project (EMP), a consortium which processed similarly public microbial 16S data into a global collection from diverse biomes, represent an unprecedented opportunity to address specific ecological questions and to unveil environmental microbiomes from diverse niches (24–26). As an example, a recent first study performed on EMP data revealed that Myxococcales prefer non-saline soil and sediments, followed by saline environments, but rarely appear in host-associated environments (26). Thus, we, herein, focused our investigation especially on non-saline soil data from EMP consortium to address the ecological diversity and relationships of Myxococcota in these ecosystems. The aims of this work were to (i) assess the ecological position and diversity of Myxococcota within microbial communities in diverse soil environments and (ii) examine potential associations between Myxococcota and other microorganisms in field soils. To further evaluate the importance of these associations, we compared them with those observed in an artificial environment where Myxococcota are not well established; this comparison revealed far fewer interactions. Overall, our analysis suggests that Myococcota may act as one of the most central ecological drivers in field soils.

## RESULTS

### Abundance of Myxococcota communities in soil environments

We started by investigating the presence of Myxococcota among microbial communities across 13 non-saline soil environments, classified by the EMP ontology (EMPO). The phylum Myxococcota was present in all these diverse habitats and distributed worldwide (Fig. 1). Myxococcota co-occurred among 6 to 15 microbial phyla, with an abundance ranging from 6.3% to 0.8% in these habitats. These results are consistent with published studies showing that Myxococcota are ubiquitous in soils, although their abundance can be quite variable.

**Figure 1.**
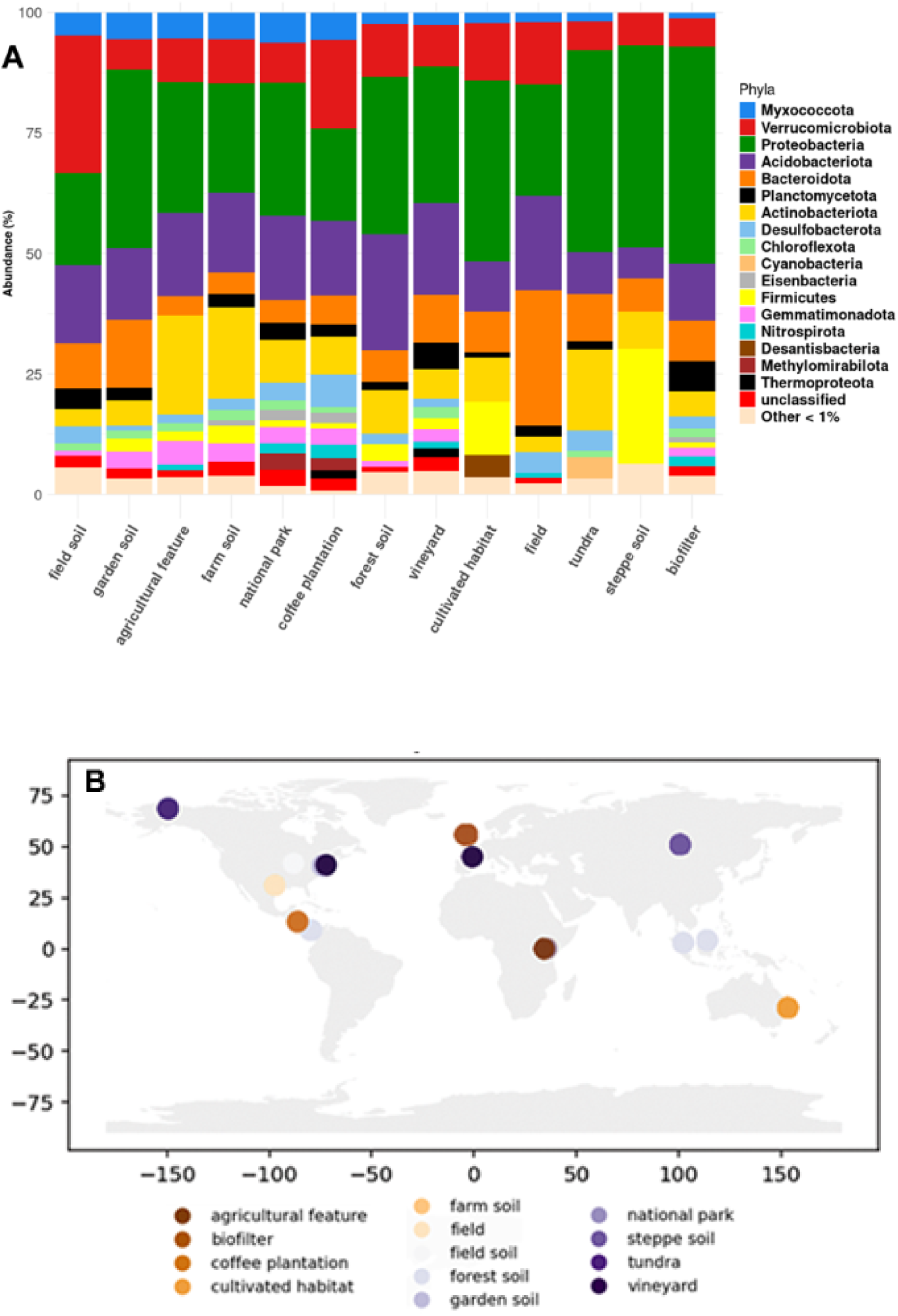
Relative abundance of Myxococcota among microbial communities in 13 soil environments (Panel A). Geographical distribution of these soil habitats across the planet (Panel B).

To further elucidate how Myxococcota interact with soil microbial communities, we examined the α/β-diversity, differential abundance and co-occurrence networks of microbial lineages between two habitats selected from the 13 soil environmental niches: the field soil and biofilter, respectively. We chose these environments for two reasons: First, they were the richest datasets in terms of samples, each represented by 690 samples – allowing robust statistical analyses - and showed distinct abundances and ranks of Myxococcota (4.9% vs 1.3%, ranks 5 vs 11, respectively) as well as differences in microbial community patterns dominated by Verrucomicrobiota (28%) and Proteobacteria (45%), respectively (Fig. 1A). Second, this choice allowed comparison most importantly, *i.e.,* the comparison of Myxococcota in communities found in between a natural (field soil (a grassland planted with switchgrass) and an artificial habitat (sand biofilter). Specifically, sand biofilters are water treatment systems that clean water by passing it slowly through layers of sand where microorganisms living in the sand remove contaminants, combining physical filtration and biological treatment.

### Diversity of microbial communities in field soil and sand biofilter environments

We first examined both α/β-diversity of microbial and Myxococcota communities in each of these environments. The α-diversity indices (richness and Shannon) of microbial communities in field soil were significantly larger than in sand biofilter (Fig. 2A, Kruskal-Wallis tests, *p-adjust* = 2.2e^-16^). Moreover, using Venn diagram analysis, we found that there were more taxa present exclusively in field soil (62.7%) than in sand biofilter (29.4%) microbial communities, and only 7.9% of taxa formed the core community between both habitats (Fig. S1A). Similarly, α-diversity analysis of Myxococcota community was also significantly higher in field soil than in sand biofilter (Fig. 2B, Wilcoxon tests, *p-adjusts* = 6.8e^208^ and 5.8e^-166^, resp.). Moreover, there were more unique Myxococcota taxa in field soil (73.5%) than in sand biofilter (18.3%). However, only 8.2% represented the core Myxococcota community between both habitats (Fig. S1B). Examination of the taxonomic composition and abundance of these taxa at the order level revealed that Myxococcota communities in both field soil and sand biofilter habitats were dominated by Polyangiales (56.5% & 37.4%, resp.) and then Haliangiales (19.4% & 14%, resp.), followed by Myxococcales and Palsa-1104 (8% & 6.6%, resp.) in field soil or UBA9160 and Myxococcales (13.5% & 11.8%, resp.) in sand biofilter (Fig. 2C).

**Figure 2.**
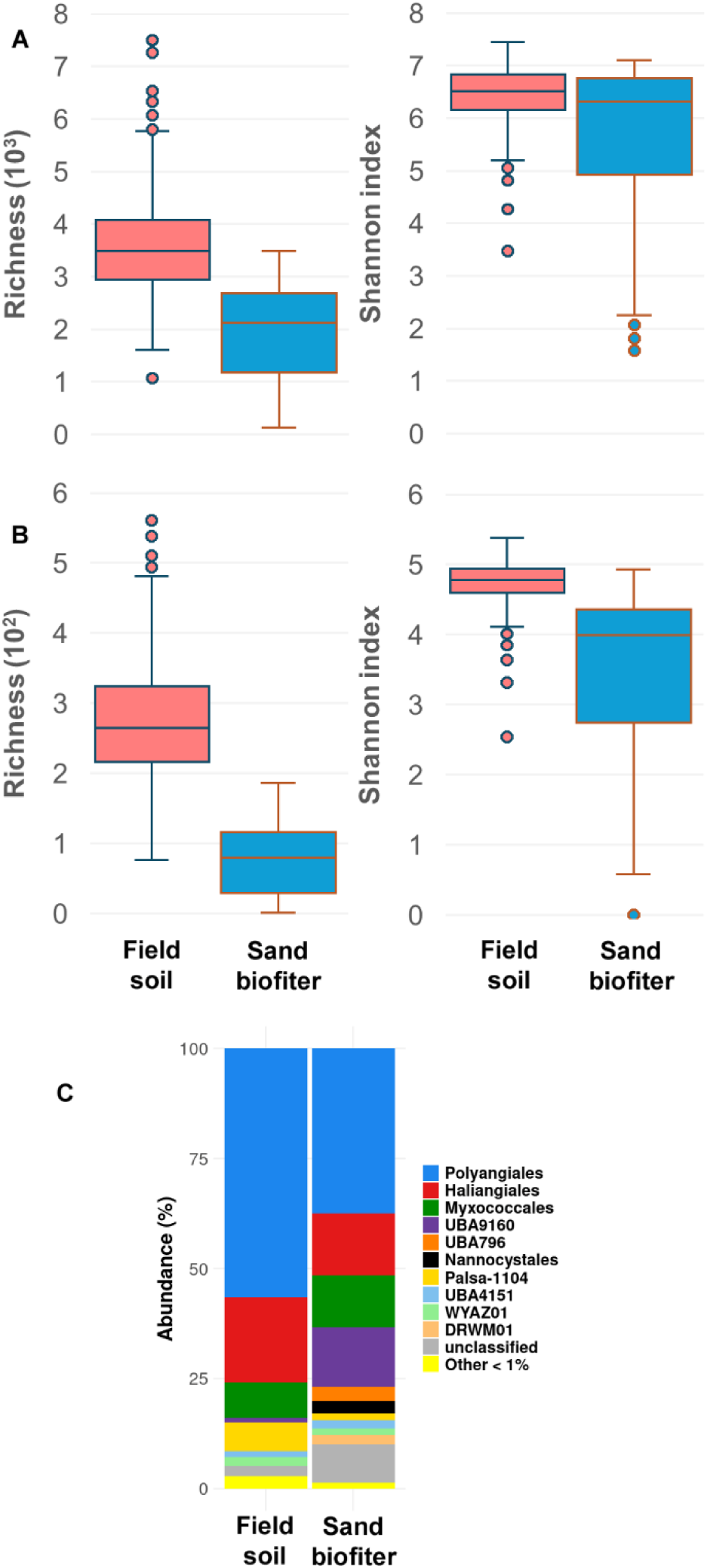
Alpha diversity of microbial and Myxococcota communities (Panels A and B, resp.) in soil habitats. Panel C. Diversity and abundance of Myxococcota communities at the order level in soil environments.

To test how diverse microbial communities are in each habitat, we used beta diversity analysis which measures the differences in community composition between sites of each habitat. Principal coordinate analysis (PCoA) based on beta diversity distances indicated a significant separation (PERMANOVA, *P(>F)* = 0.001) between field soil and biofilter samples mainly along the first axis (35.9% of the explained variance) (Fig. 3). These beta diversity differences were also supported by the betadisper analysis that showed significant differences (P*(>F)* = 2.2^e-16^) between field soil (average distance to median = 0.408) and biofilter (0.475) samples. Overall, field soil samples clustered closely in ordination space and exhibited a lower dispersion, suggesting the possibility of similar community diversity across these samples. In contrast, biofilter samples were more widely distributed and showed higher dispersion, indicating greater variability in microbial community structure. The curved distribution of biofilter samples in the ordination may be consistent with a gradient in community composition, although the drivers of this pattern cannot be determined from the ordination alone.

**Figure 3.**
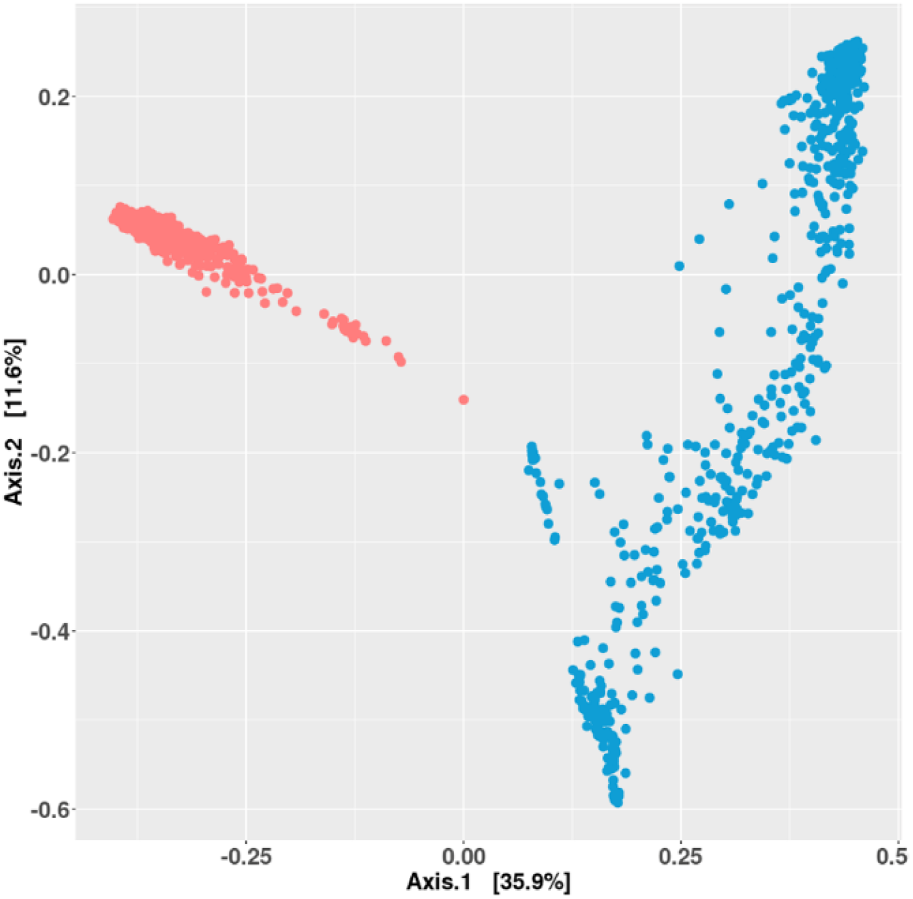
Beta diversity of microbial communities in field soil (in salmon pink) and sand biofilter (in turquoise) environments as inferred using Bray-Curtis dissimilarity and Principal Coordinates Analysis (PCoA).

Last, we found 19 microbial phyla displaying significant differences in the abundance between field soil and sand biofilter (Fig. 4, LDA score >=3.3, *p-value* <= 0.004). Remarkably, the Myxococcota was significantly more prevalent in field soil together with 8 other microbial lineages including Verrucomicrobiota, Acidobacteriota, Bacteroidota, Desulfobacterota, Bacteria_unclassified, UBA10199, Elusimicrobiota and Patescibacteria. In contrast, 10 phyla were more represented in the sand biofilter, among them the Bdellovibrionota, regrouping many other bacterial predator species, along with Proteobacteria, Firmicutes, Actinobacteriota, Planctomycetota, Nitrospirota, Gemmatimonadota, Thermoproteota, Eisenbacteria and Cyanobacteria.

**Figure 4.**
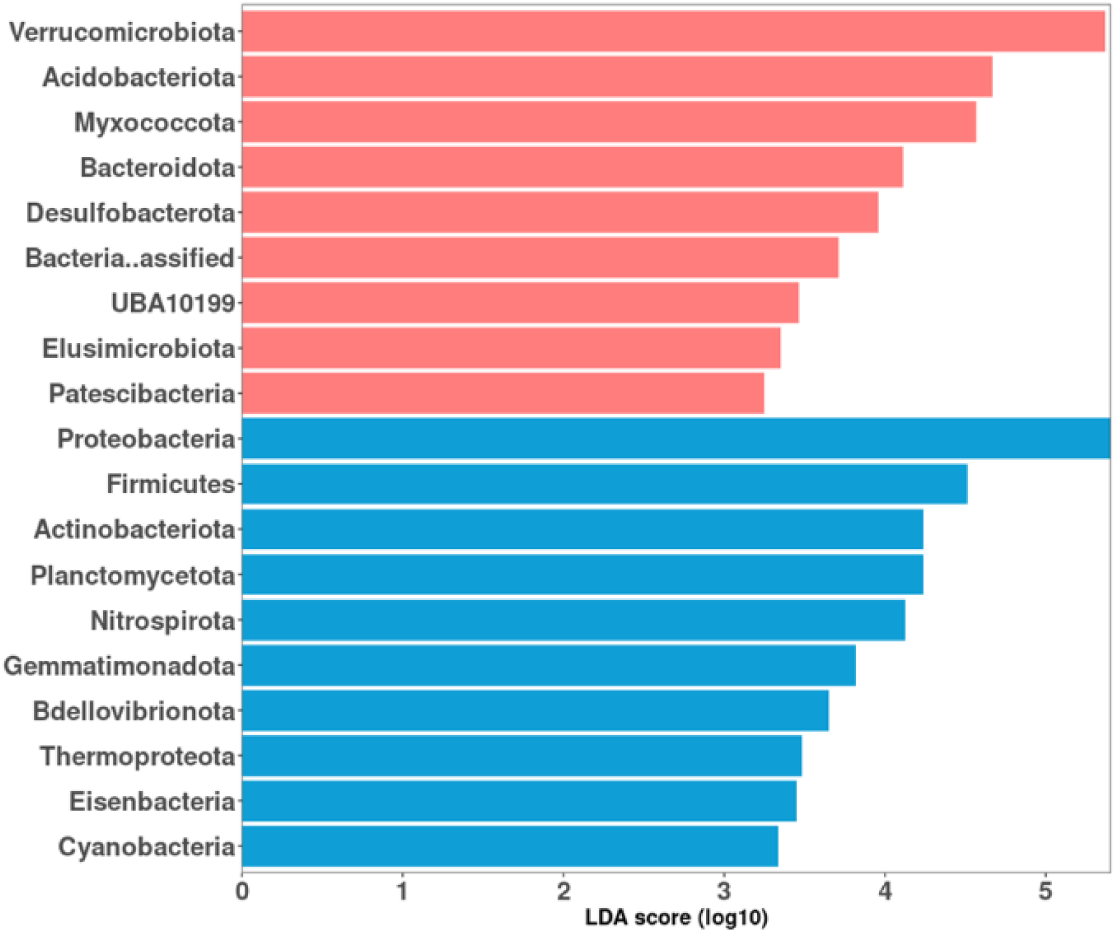
Differential abundance of microbial phyla between field soil (in salmon pink) and sand biofilter (in turquoise) environments, ranked by effect size using LEfSe analysis.

### Structure of the field soil microbial community

We next explored the structure of the two habitats, starting from the field soil where the high relative abundance of Myxococcota (4.9%) suggests that they are an important component of the microbial community. Therefore, we first constructed the field soil co-occurrence network based on Spearman’s correlations to elucidate the structure of this community (Fig. 5A). This network comprised 684 nodes (or taxa) connected by 13,624 significant correlations (|ρ| ≥ 0.75) (Table 1). Using this statistical cut-off, the network was strongly dominated by positive associations, with 99.9% of the edges reflecting positive correlations, suggesting that most microbial taxa tended to co-vary in abundance across samples. This prevalence of positive correlations may suggest shared environmental preferences, co-occurrence in similar niches and/or potential beneficial interactions (e.g., mutualistic) among taxa within the field soil microbial community. In contrast, negative correlations were rare, accounting for only 0.1% of the edges. This should be interpreted with caution because the chosen methodology might also limit their detection due to the compositional nature of the data (not absolute counts) and statistical filtering (negatives are harder to detect and tend to disappear after multiple corrections (27).

**Figure 5.**
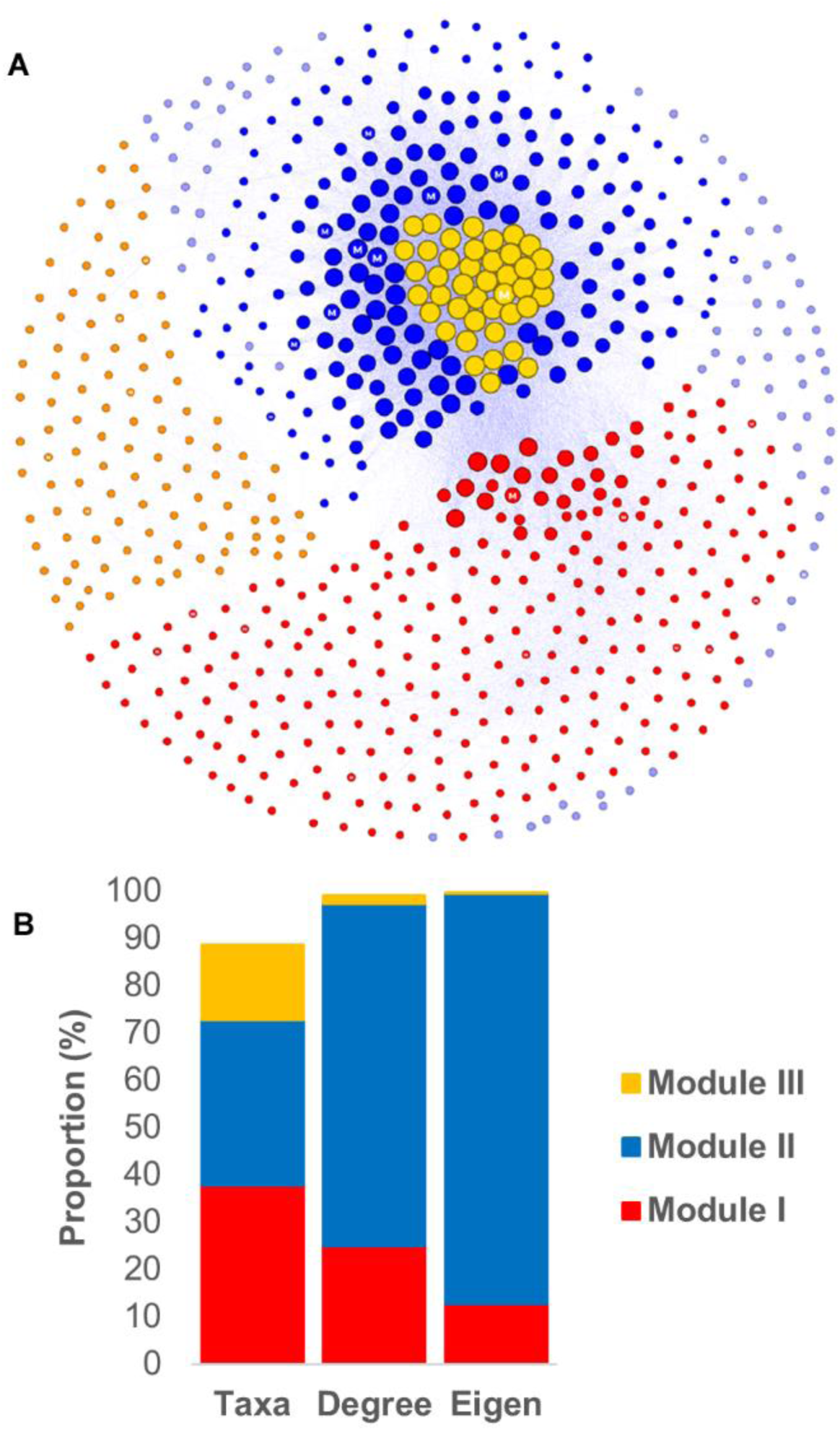
Network of co-occurring microbial taxa in field soil environment (**A**). Nodes (or taxa) of three major modules (I, II and III) are colored in red, blue and orange, respectively. Nodes of minor groups are colored in blue-magenta. Node size depicts eigen centrality value. Hub nodes are indicated by gold color. Myxococcota’s nodes are indicated with the letter M. Edges with positive and negative correlations are colored in blue-magenta and lightSalmon, respectively. **B**, Global distribution of microbial taxa and their degree and eigen centralities across three major modules in field soil network.

**Table 1.** Global topological features of soil microbial networks.

| Features | Field soil | Sand biofilter |
| --- | --- | --- |
| Nodes | 684 | 926 |
| Edges | 13,624 | 92,146 |
| Positive edges | 13,608 (99.9%) | 85,600 (92.9%) |
| Negative edges | 16 (0.1%) | 6,546 (7.1%) |
| Nodes positively correlated | 683 (99.9%) | 926 (100%) |
| Nodes negatively correlated | 18 (2.6%) | 476 (51.4%) |
| Modularity | 0.271 | 0.107 |
| Modules | 28 | 14 |
| Average degree | 39.84 | 199.02 |
| Clustering coefficient | 0.679 | 0.788 |

Nevertheless, the network exhibited a strong modular structure (modularity = 0.271), clustering taxa into three main modules (I, II, and III), potentially reflecting distinct functional or ecological assemblages, along with 25 minor, peripheral or isolated groups (Fig. 5A, Table 1). The distribution of microbial taxa was highest in modules I (37.9%) and II (34.8%), compared to module III (15.8%) (Fig. 5B). However, module II clearly emerged as the structural core of the network: it harbored 72.4% of all degree centrality and 86.7% of all eigenvector centrality (Fig. 5B), indicating that taxa in this module were both highly connected and influential in the overall network topology. In contrast, modules I and III exhibited lower connectivity and centrality (Fig. 5B), suggesting more peripheral or specialized roles in the field microbial community. This underscores the ecological significance of module II, potentially acting as a hub for key microbial interactions and functions in field soil ecosystems.

### Position of the Myxococcota in the field soil microbial community

To better understand the ecological role of the Myxococcota within the field soil microbiome, we first examined their distribution and properties across the three main modules of the co-occurrence network (Fig. S2). Thirty Myxococcota taxa were identified: 11 (1.6%) in each of modules I and II, and lower in module III (5, 0.7%) and in minor groups (3, 0.4%). However, the Myxococcota were significantly more connected in module II, displaying higher degree and eigen centralities (3.6% and 4.5%, resp.) than in modules I (0.9% and 0.5%, resp.) and III (0.1 and 0.01%, resp.). In module II, the Myxococcota was found among the most connected phyla, ranking amongst the most highly connected phyla in terms of degree and Eigen centrality, ranking at the 6^th^ position after Acidobacteriota, Proteobacteria, Verrucomicrobiota, Bacteroidota and Planctomycetota (Fig. S2). Moreover, it outperformed several phyla such as the Firmicutes and Bdellovibrionota, the other main predatory phyla which displayed very low centralities (<=0.01%) in module II (Fig; S2).

We next examined the sub-network of Myxococcota interactions within each of the modules. In total, the 27 Myxococcota taxa were positively connected to 294 taxa by 1,208 edges, across all three modules, but their connectivity patterns and associations varied markedly by module (Figs. 6 and S3). Module II emerged once again as a key ecological unit: the 11 Myxococcota taxa in this module exhibited the highest degree of connectivity, forming 78.6% of their links with neighboring taxa, a large majority of them within module II. These connections spanned 16 microbial phyla, with dominant associations to Proteobacteria (16%), Acidobacteria (14.9%), Verrucomicrobiota (12.6%) and Bacteroidota (9.8%), indicating that Myxococcota in module II are deeply embedded in a diverse and functionally rich microbial context.

**Figure 6.**
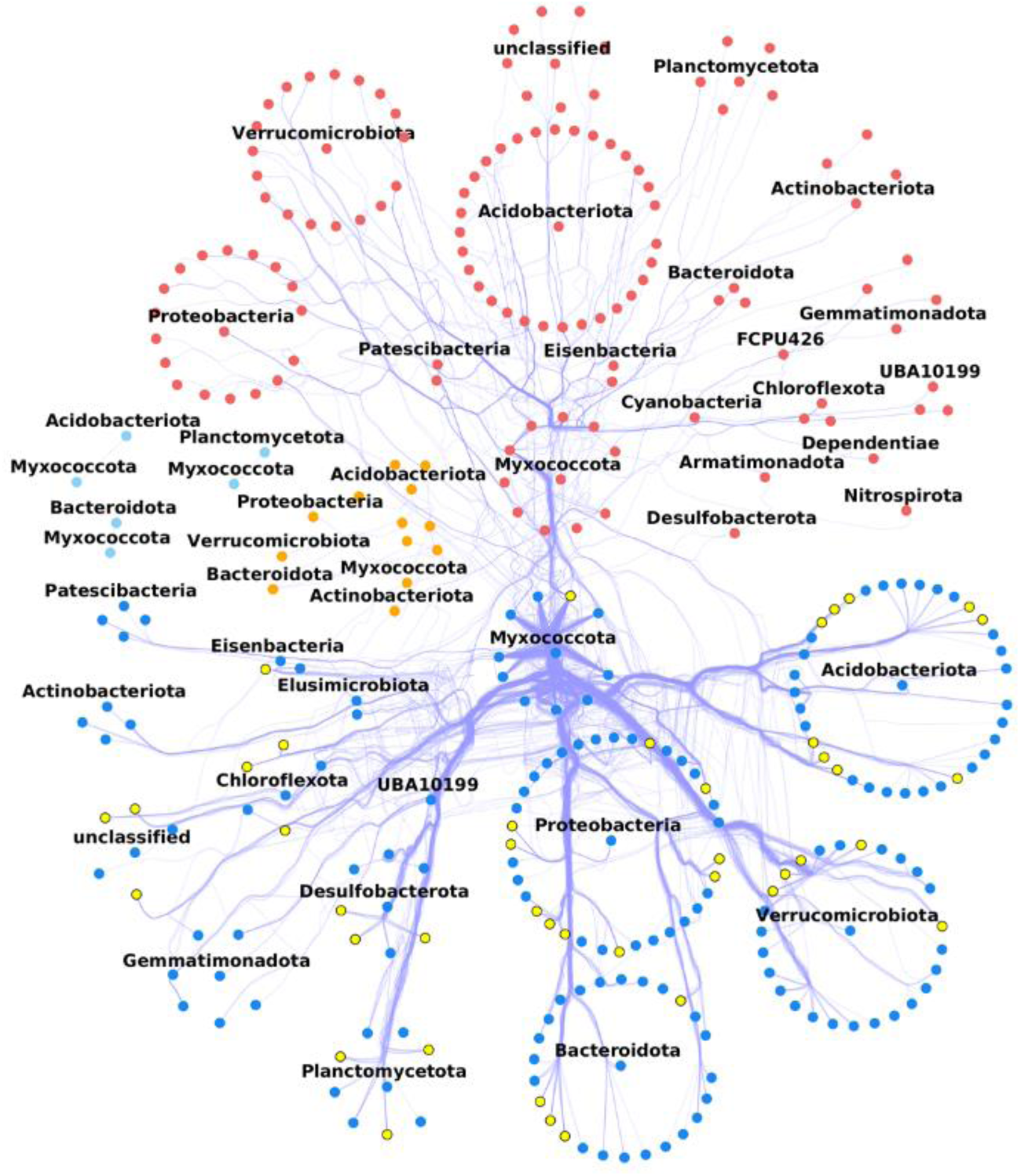
Subnetwork exhibiting Myxococcota neighbors (or taxa) according to their corresponding three major modules (I, II and III) in the field soil environment. Neighbors are grouped by their corresponding phylum. Nodes (or neighbors) of modules (I, II and III) are colored in red, blue and orange, respectively. Nodes of minor groups are colored in blue-magenta. Hub taxa of the network are represented by nodes colored in yellow and black border. Edges connecting nodes are bundled and depict positive (blue-magenta) and/or negative (lightSalmon) co-occurrence correlation.

By contrast, the 11 Myxococcota taxa in module I had significantly fewer associations (20% connectivity, mainly with taxa of this module), despite linking to a broader set of 19 phyla, most of which (i.e., 16) overlapped with those in module II. Three additional phyla—Dependentia, Nitrospirota, and FCPU426—were uniquely connected to Myxococcota in module I, suggesting more specialized or peripheral interactions.

The 5 Myxococcota taxa in module III showed the lowest connectivity, with only 1.2% of their possible links realized, and associations limited to 6 phyla, all of which were also present in the other two modules.

Overall, these patterns reinforce the idea that module II serves as the central ecological hub in the field soil microbial community, with Myxococcota acting as key connectors in this module. Their broad and dense associations with multiple microbial phyla—especially in module II—suggests that they deeply influence microbial community dynamics in the soil.

### Comparison with an engineered environment: sand biofilter

To get a deeper understanding of the significance of the interactions observed in the field soil habitat, we further characterized the sand biofilter for a comparison with an artificial habitat. Indeedsand biofilters are porous media used for water treatment and thus their composition is expected to fluctuate and host a specialized microbial community (for example a biofilm) adapted to rapid processing and resource competition, very distinct from ecology that would be associated to a soil environment.

In this environment, we showed that Myxococcota displayed a much lower abundance (1.3%). Interestingly, the sand biofilter microbial network exhibited a much higher complexity and connectivity. It consisted of 926 taxa connected by an extensive 92,146 associations, resulting in a significantly higher edge density (0.233) and average degree (199.02) (Fig. 7A, Table 1). While positive associations still dominated (92.9% of edges), the presence of negative connections was much more pronounced than in field soil and this despite the positive bias introduced by correlations analysis: 51.4% of taxa were involved in some negative associations, representing 7.1% of total edges. This suggests a more competitive or dynamic microbial environment within the engineered sand biofilter, likely driven by the constant environment changes linked to water treatment.

**Figure 7.**
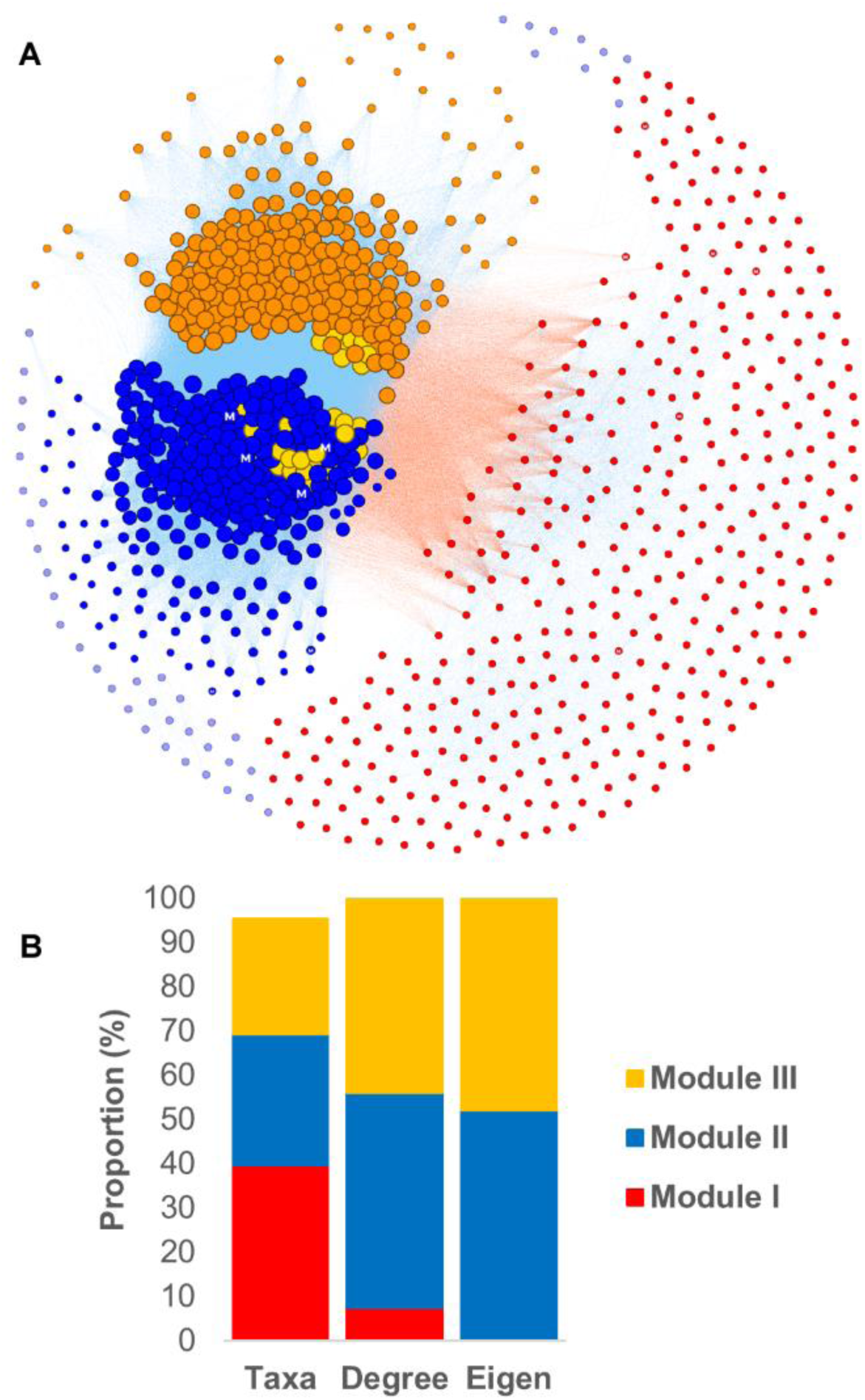
Network of co-occurring microbial taxa in sand biofilter habitat (**A**). Nodes (or taxa) of three major modules (I, II and III) are colored in red, blue and orange, respectively. Nodes of minor groups are colored in blue-magenta. Node size depicts eigen centrality value. Hub taxa are indicated by black border color of nodes. Myxococcota’s nodes are indicated with the letter M. Edges with positive and negative correlations are colored in blue-magenta and lightSalmon, respectively. **B**, Global distribution of microbial taxa and their degree and eigen centralities across three major modules in sand biofilter network.

The sand biofilter microbiome was less modular (0.107 *vs* 0.201) and organized into three major modules (I, II, and III) and only 11 minor clusters (Fig. 7A, Table 1). Module I contained the highest proportion of grouped taxa (39.7%), followed by two closely related modules II and III (29.6% and 26.0%, respectively) (Fig. 7B). However, when examining microbial centralities, both modules accounted for the majority of the network’s central associations—48.6% and 43.9% for module II, and 51.7% and 41.9% for module III—compared to only 7.4% and 0.4% in module I (Fig. 7B).

Notably, the high proportion of negatively associated taxa (51.4%) observed in the sand biofilter network was predominantly found between module I and modules II/III (Fig. 7A, Table 1), indicating strong negative interactions. This pattern suggests the presence of functionally distinct and potentially competing/transient microbial communities, consistent with the competitive and resource-driven nature of the biofilter environment.

### Position of the Myxococcota in the sand biofilter microbial community

In the sand biofilter network, we detected 12 Myxococcota taxa separated across modules as follows (Figs. 7A/B and S4): 6 (0.65%) and 6 (0.65%) taxa in modules I and II, but none (0%) in module III, respectively. However, these Myxococcota displayed higher degree and eigen centralities in module II (1% and 1.1%, resp.) than in module I (0.07% and 0.02%, resp.) (Fig. S4). Interestingly, in module II, the centrality patterns changed greatly compared to module II in the field soil network. This module was mostly structured by Proteobacteria, Bacteroidota and to a lower extent by Verrucomicrobiota, Firmicutes, Bdellovibrionota, Planctomycetota, Acidobacteriota and Actionobacteriota (Fig. S4). Myxococcota were less central than all these taxa even ranking after Firmicutes and Bdellovibrionota, contrarily to what was observed in field soils (Fig. S4). Interestingly, the centrality of Acidobacteriota also decreased drastically in the sand biofilter modules in comparison to the field soil modules (Figs. S2 and S4, resp.). Analysing Myxococcota interactions more closely in the sand biofilter sub-network, we observed notable differences in the number, composition and profile of microbial neighbors associated with Myxococcota (Fig. 8) compared to the field soil subnetwork. Within this sub-network, the 12 Myxococcota taxa were distributed between modules II (6 taxa) and I (6 taxa). However, the number and type of correlations were completely distinct for Myxococcota in these modules (Figs. 8 and S5). In module II, the Myxococcota were highly interconnected, accounting for 93.3% of all the Myxococcota links in the entire. They were also mainly associated with Proteobacteria (45.5%) and Bacteroidota (16.9%), along with links to Verrucomicrobiota, Planctomycetota, Acidobacteria, Gemmatimonadota and Actinobacteriota. Importantly significant correlations were observed with Firmicutes and Bdellovibrionota, which were absent in the field soil sub-network. In contrast, Myxococcota in module I displayed a reduced and narrower spectrum of correlations (6.7%), lacking several connections observed in module II (Firmicutes, Bdellovibrionota, Armatimonadota, Bacteroidota, Cyanobacteria, Dependentiae, Chlamydiota, and Thermoproteota), the remaining links being much weaker, contributing less than 2.7% of the correlations of the entire subnetwork.

**Figure 8.**
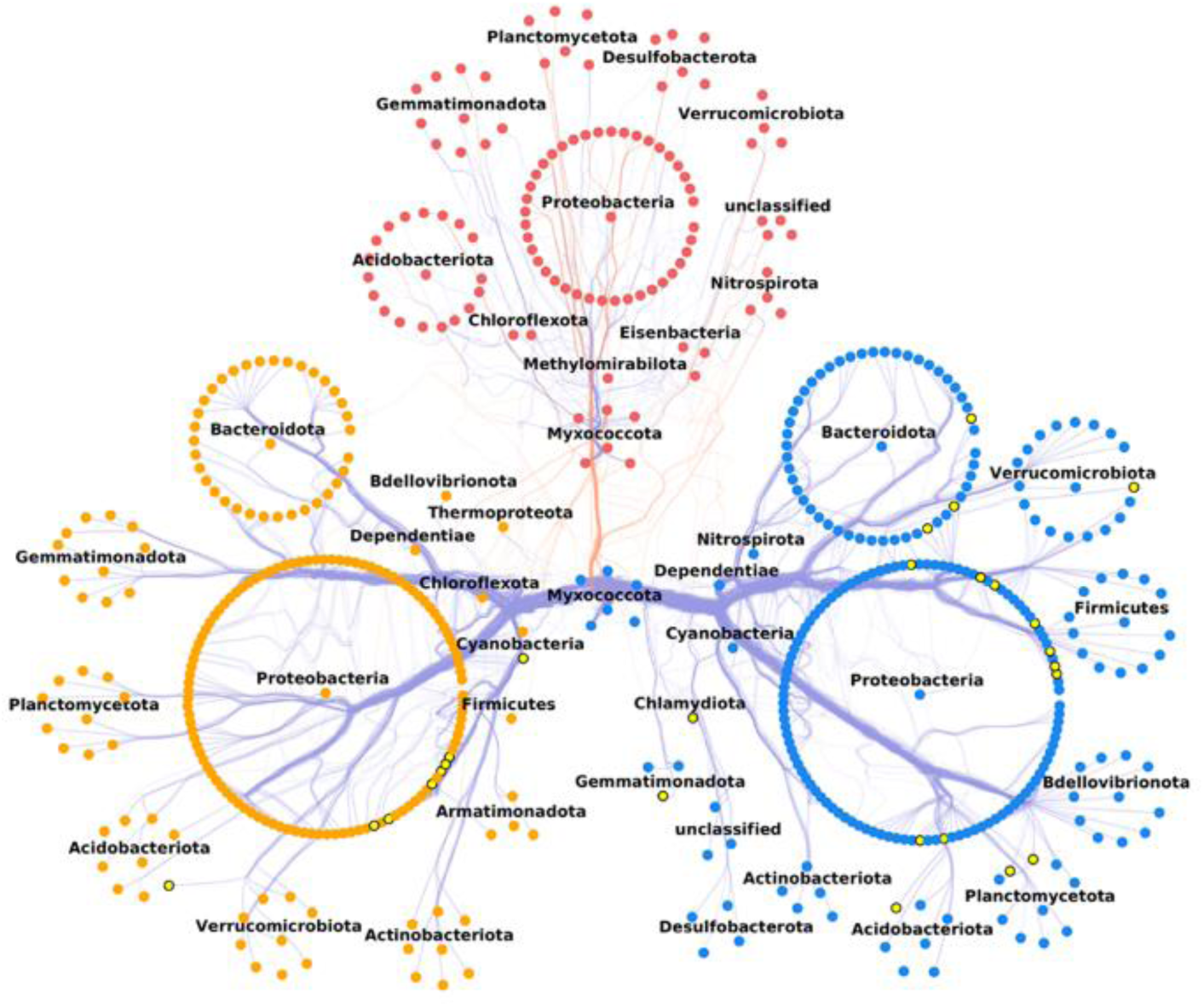
Subnetwork showing Myxococcota neighbors (or taxa) according to their corresponding three major modules (I, II and III) in the sand biofilter environment. Neighbors are grouped by their corresponding phylum. Nodes (or neighbors) of modules (I, II and III) are colored in red, blue and orange, respectively. Nodes of minor groups are colored in blue-magenta. Hub taxa of the network are represented by nodes colored in yellow and black border. Edges connecting nodes are bundled and depict positive (blue-magenta) and/or negative (lightSalmon) co-occurrence correlation.

Importantly, negative associations were observed between Myxococcota and several taxa in modules I and III (Proteobacteria, Acidobacteria, Planctomycetota, Verrucomicrobiota, Bacteroidota, Actinobacteriota, Nitrospirota, Desulfobacterota, Gemmatimonadota, Methylomirabilota), suggesting the presence of negative interactions present in the sand biofilter habitat but absent from the field soil sub-network.

### Myxococcota are central to the structure of the field soil microbial community

Altogether, the results above indicate that Myxococcota engage in a large number of potential interactions in field soil environments, suggesting they may play a central role in shaping community structure in this habitat. In contrast, this appears to be much less the case in sand biofilters, where complex communities are observed but Myxococcota are less central and more often involved in negative interactions. To explore this further, we examined the connection between Myxococcota and hub taxa within the field soil and sand biofilter networks. In the field soil network, we identified 42 hub taxa with eigenvector centrality values ranging from 0.84 to 1—above the 99th percentile of empirical distribution (Figs. 5A) (Peschel et al., 2021). By comparison, only 26 such taxa were identified in the sand biofilter network (Figs. 7A). This confirms that the field soil community is indeed more structured than the sand biofilter community.

We then examined how Myxococcota connected to hub taxa and whether they could themselves be hub taxa, or both in sub-networks extracted from networks. In the field soil sub-networks, we observed a dense connectivity revealing that Myxococcota are highly connected to hub taxa (Figs. 6 and 9A). This was specifically true for module II, where 8 out of 11 Myxococcota taxa [eigen from 0.29 to 0.79, by comparison only 2 of 11 Myxococcota taxa in module I were connected (from 0.09 to 0.52) to hub taxa], were strongly associated with hub taxa. Importantly, one Myxococcota taxon from the order Polyangiales even stood out as a hub taxon (eigen = 0.99) (Fig. 9A).

**Figure 9.**
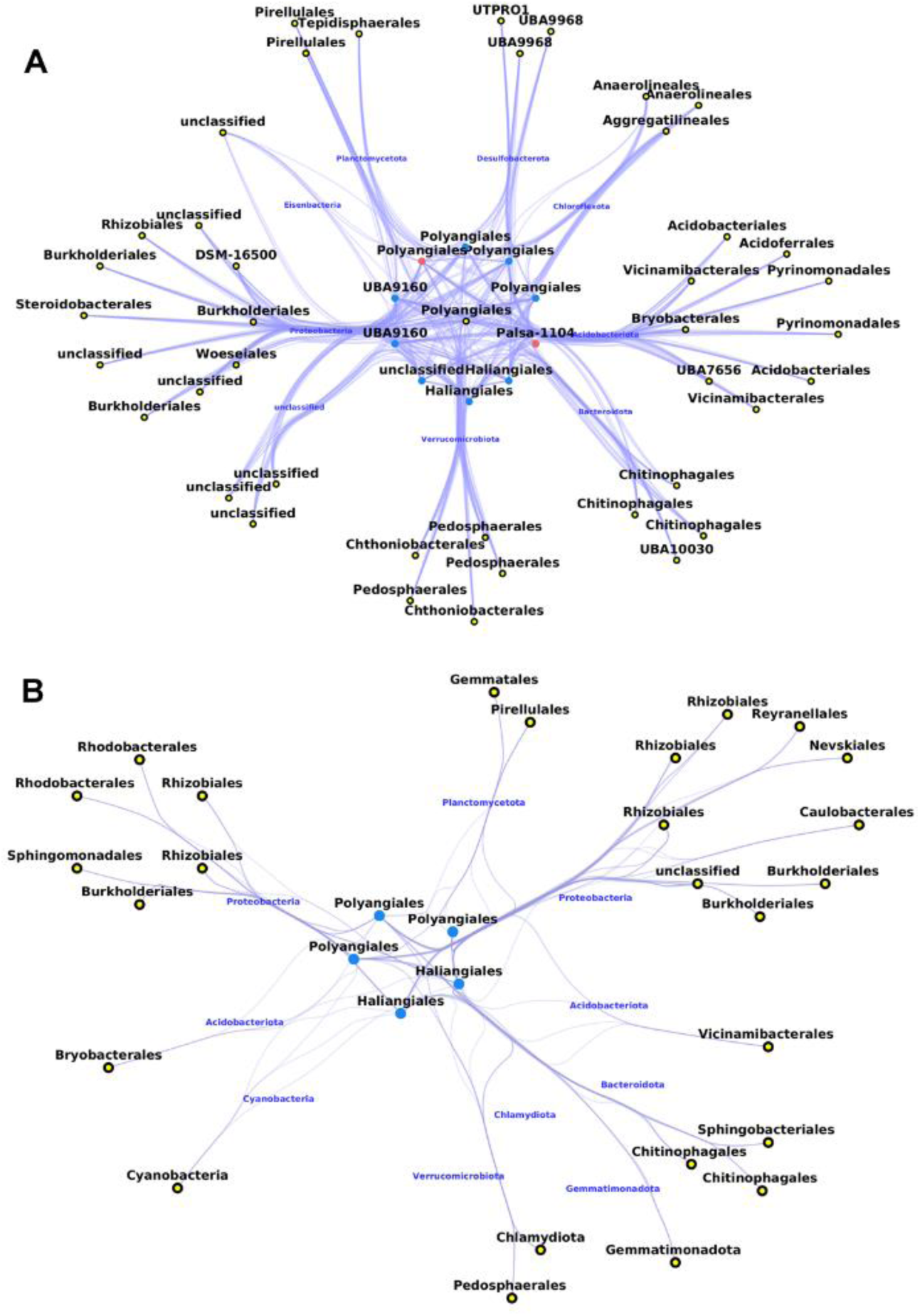
Myxococcota taxa connected to microbial hub taxa in field soil (**A**) and sand biofilter (**B**) subnetworks. Taxa are grouped and named by their corresponding phylum and orders. For field soil: the single Myxococcota hub (in yellow with black border) is from module II; the remaining Myxococcota are from modules II (in blue) and I (in red), respectively; the hub taxa of other phyla (in yellow with black border) are from module II. For sand biofilter: Myxococcota taxa are from module II (in blue) and, hub taxa of other phyla (in yellow with black border) are from modules II (in the right) and III (in the left). Edges connecting taxa are bundled and depict positive associations.

In the sand biofilter sub-networks (Figs. 8 and 9B), the connections exhibit a more diffuse and simplified architecture. But nevertheless, 5 out of 6 Myxococcota taxa in module II (eigen from 0.11 to 0.96) also showed associations with hub taxa suggesting that they also participate in the organization of the community. However, given the overall community is much less structured in the sand biofilter system.

## DISCUSSION

Soil microbiomes are central to the sustainability of multiple ecosystem functions, organic matter composition, nutrient cycling and plant production. Recent metagenomics studies suggest that bacterial predators are more common in soil microbiomes than originally suspected, raising the possibility that predator-prey interactions are important for the balance of soil ecosystems (26, 28).

To investigate this possibility, we analyzed microbiomes from various EMP non-saline soils, focusing first on the presence and abundance of Myxococcota, a bacterial phylum known to comprise major predatory taxa (Although it cannot be ascertained that all members of the Myxococcota are predatory). Our results show that Myxococcota are indeed present in all the examined non-saline soil microbial habitats, confirming both their dispersal capacity and the ubiquitous nature of this lineage. Nevertheless, the relative abundance of Myxococcota varied with soil type, ranging from soils with higher abundance (4.9–6.3% of the detected taxa) to those with lower abundance (0.8–2.6% of the detected taxa). Further work is required to elucidate the drivers underlying this variation. It is likely that both abiotic are important because the abundance of myxobacterial communities has been shown to correlate with soil pH, carbon-to-nitrogen ratio, and temperature in grassland ecosystems (29). These factors may influence biotic interactions and shape competitive dynamics within microbial communities.

To further elucidate the ecological significance and functional role of Myxococcota communities in soil environments, we performed a comparative microbiome analysis of a natural habitat (field soil) and an artificial habitat (sand biofilter used for water treatment). This analysis included assessments of microbial diversity and co-occurrence networks, under the premise that such a comparison would reveal how Myxococcota integrates in natural soils, versus a more dynamic, artificially fluctuating environment. The two habitats differed markedly: the field soil exhibited relative homogeneity in community composition (as indicated by low β-diversity) and high overall microbial diversity (high α-diversity), whereas the sand biofilter was characterized by greater variability in microbial composition and lower diversity. The Myxococcota diversity was higher in field soil than in sand biofilter habitats, which is in agreement with another study that correlated Myxococcota diversity to bacterial diversity in other soil ecosystems (30). Interestingly, the Myxococcota communities were dominated by Polyangiales and Haliangiales, followed by the Myxococcales in both environments. This is similar to Myxococcota compositions described in other soil environments, suggesting a conserved ecological structure within this phylum (3, 31).

To get better understanding of the ecological function of Myxococcota, we next constructed co-occurrence networks describing both habitats to infer hypotheses on mechanisms of biotic interactions or shared environmental preferences among microbial taxa (32–34). The resulting co-occurrence networks were markedly distinct. The field soil microbiome was dominated by positive associations (99.9% vs 92.9%) and a high number of hub taxa (42). This structural organization is especially prominent in module II, which represents a dominant sub-community in the field soil microbiome. Overall, this network structure suggests a tightly knit community, a potential signature of a stable ecosystem.

In contrast, the sand biofilter community appears more complex, it is less cohesively integrated by positive interactions, and it contains stronger evidence of negative associations. This dynamic structure is particularly apparent in the striking negative correlations between module I and modules II and III, suggesting pronounced fluctuations and antagonisms between sub-communities. The lower number of hub taxa further suggests a lower organization of the sand biofilter microbiome. These contrasting patterns support the conclusion that the two microbiomes reflect fundamentally different habitats, prompting further investigations into the role of Myxococcota in each context.

The connections of Myxococcota in the field soil suggest that they are well integrated in soil communities, and thus potentially important players in the trophic chain. This is particularly evident in the network where Myxococcota exhibited high centralities, and one Polyangiales taxon with eigenvector equal to 0.99—thus classified here as a hub taxon. This evidence was especially strong in module II. Interestingly, in the sub-network, Myxococcota in this module shared high connections to three phyla, Acidobacteria (16%), Proteobacteria (14.9%) and Verrucomicrobiota 12.6%). In contrast, the Myxococcota were more peripheral in the sand biofilter environment (module II, 1% and 1.06%), and they also were involved in negative correlations. These negative correlations could be especially meaningful because Spearman’s correlations tend to bias for positive correlations. Moreover, Myxococcota in module II was mainly connected to Proteobacteria (45.5%) and exhibited comprehensive associations e.g., with Firmicutes (2.9%) and Bdellovibrionata (2.5%) which lacked in field soil sub-network. There are, however, several limitations that need to be considered when considering these results:

i. Positive and negative correlations do not necessarily indicate direct interactions. While mutualistic cooperative interactions may account for some positive correlations, they can also arise from taxa occupying similar ecological niches (ecological filtering).
ii. Predatory interactions cannot be predicted from this analysis. First, while the Myxococcota phylum includes well-established predatory lineages (e.g., the Myxococcales), the absence of definitive genetic markers for bacterial predation prevents us from determining whether all detected taxa are indeed predators (6,7). Second, predator-prey interactions can be reflected by both positive and negative correlations depending on the environment and these interactions can fluctuate over time (35). In ecosystems, predation is typically characterized by the coexistence and co-evolution of predator and prey, as described by the Red Queen hypothesis (36, 37), but depending on activity and the environment, predator and prey populations can fluctuate deeply by changing the connections and modules within the network (35).
iii. We are using data from the Earth Microbiome Project, which provides curated and standardized 16S datasets allowing site comparisons. However the analysis remains geographically limited, the field soils samples were all collected in the USA and the biofilter in Scotland, which could introduce a bias and but nevertheless provides a first snapshot of Myxococcota ecology.

Despite these limitations, our findings reveal clear pattern differences between these habitats and suggest that Myxococcota interactions are more deterministic in field soils (versus stochastic in biofilter), suggesting ecological integration. This pattern seems to agree with a recent study which showed that myxobacteria community assembly is dominated by deterministic processes in agricultural soils (38). Defining the mechanisms will require deeper analyses in natural as well as in laboratory contexts including the development of synthetic communities. Further metagenomic studies in the field are required to clearly identify the types and dynamics of biotic interactions at the species level and thus potentially uncover direct interactions between organisms. In addition, the precise geochemical determination of the spatial environment will be needed to determine the abiotic components in which these bacteria thrive. Both approaches will be important to discriminate direct interactions from indirect effects such as independent influence of the environment. This will be extremely challenging given the complexity of soil ecosystems in terms of bacterial species, but also other organisms including fungi, protozoa and nematods. However, the goal is worth pursuing, as the ability to assess soil health based on predator–prey network structure could not only provide a diagnostic tool for soil condition but also open avenues for restoration through the reintroduction of key predators or prey.

## MATERIALS AND METHODS

### EMP 16S data and metadata

The 16S metagenomic data and metadata were from obtained from EMP website (https://earthmicrobiome.org/). This consortium is a massively collaborative effort to characterize microbial life on this planet which has curated and standardized 16S data and metadata using unified methodologies including collecting, (pre)processing, ontologies and curating 16S metabarcoding data and metadata from diverse microbial community studies across the globe. We extracted samples at the level 3 of the EMP ontology [empo_3 = soil (non-saline)] from Deblur 16S-V4 region. Taxonomic assignment of Amplicon Sequence Variants (ASVs) was updated using a Bayesian classifier trained with Greengene2 database Version 2022.10 using QIIME2 (39). We obtained a metadata frame and a BIOM matrix containing 2,514 soil samples and 110,897 features (or ASVs or taxa) and a total read counts 85,798,519 reads.

We used less stringent filters to remove taxa exhibiting an abundance of less than 3 reads / sample and a prevalence in less than 10 samples. Subsequent analyses were then performed on 39,909 ASVs present in 2,514 samples and with 97% of the total abundance (*i.e.,* of 85,798,519 read counts). These samples were classified into 13 environmental categories based on the EMP ontology feature and material, collected from diverse countries, as follows: field soil (n = 696 samples, USA), sand biofilter (n=816, Scotland), cultivated habitat (n= 214, Australia), steppe soil (n=204, Mongolia), garden soil (n=145, USA), forest soil (n=101, Malaysia, Mongolia and Panama), vineyard (n=65, USA), field (n=62, USA), coffee plantation (n=60, Nicaragua), agricultural feature (n=54, Kenya), farm soil (n=47, Kenya), tundra (n=28, USA) and national park (n=22, Kenya). All data extraction and analysis were performed using QIIME2 algorithms (40, 41) and Phyloseq packages under R/Studio interface (42).

### Community diversity analysis and differential abundance

Alpha diversity of microbial taxa for each of the field soil and biofilter environments was estimated, without rarefaction of read counts, using richness and Shannon’s entropy indexes. For each habitat, we randomly extracted 690 samples from the original dataset and filter rare taxa (with abundance < 3 reads/sample and prevalence less than 10 samples). Differences in alpha diversity indexes between habitats were evaluated using Wilcoxon and Kruskal-Wallis tests. The beta diversity between habitats was explored from normalized datasets using the Bray-Curtis dissimilarity index and PCoA ordination method. Statistically significant differences in beta diversity dissimilarities between both soil habitats were statistically assessed by adonis2 (PERMANOVA) and, betadisper [for homogeneity of groups dispersion (Variances)] and ANOVA, functions implemented in the Phyloseq and the Vegan packages using RStudio interface, respectively.

To investigate whether Myxococcota phylum were either differentially abundant between two habitats, we calculated the LEfSe (LDA effect size method) and then performed a comparative network analysis. The LEfSe analysis integrates statistical significance with biological consistency (effect size) estimation of differentially abundant phyla. It is based on the Kruskal-Wallis test (<=0.01) to detect features with significant differential abundance with respect to the class of interest and linear discriminant analysis (LDA score >= 3.3, 43) to estimate the effect size of each differentially abundant feature and implemented in the MicrobiomeMarker R package (44, 45).

### Co-occurence network

Co-occurrence networks are used to infer potential ecological associations among microbial taxa in soil metagenomic datasets (32, 33, 46, 47). In these graphical representations, nodes represent ASVs (or their microbial taxa), and edges (connections between nodes) represent significant correlations in their abundance profiles across multiple soil samples. (33, 48, 49). This approach helps to uncover potential microbial interactions between taxa or shared environmental preferences (32, 34).

We constructed co-occurrence networks of both field soil (4,159 taxa) and biofilter (2,524 taxa) habitats after removing taxa with less than a total of 10^3^ reads in 690 samples to reduce the high dimensionality of our data, which may be the source of false-positive predicted associations. These networks were performed using Spearman’s correlation (|r| ≥ 0.75) on clr-normalized counts which aimed to avoid compositional effects (33, 48–50). Network clustering was performed using fast_greedy optimization algorithm implemented into the NetCoMI R package (33). Network visualization and calculation of a local topological property and taxa connectedness of networks and subnetworks (i.e., number of nodes and edges, degree and eigen centralities) as well as the prevalence (as total number of taxa) of taxa and phyla were performed using NetCoMI and Cytoscape software (33, 51). To detect potential hub taxa of each habitat, we used high eigenvector centrality (at a cutoff value above the empirical 0.99% percentile), a metric detecting the most central nodes while considering the relevance of their neighbors in the co-occurrence networks (33, 52).

## AUTHOR CONTRIBUTIONS

KE and TM conceived the project. KE performed bioinformatic analysis. KE and TM wrote the manuscript. BP helped with the interpretation of the results.

## CONFLICTS OF INTEREST

The authors certify that they have no conflicts of interests in relation to this research.

## ACKNOWLEDGMENTS

The study was funded by CNRS and AIX-MARSEILLE university funds to TM

## SUPPLEMENTAL MATERIAL

Supplemental material is available online only.

**Supplemental file 1**, PDF file.

## DATA AVAILABILITY

The EMP (Earth Microbial Project) dataset is available at ftp://ftp.microbio.me/emp/release1.

## Notes

### Competing Interest Statement

The authors have declared no competing interest.

## REFERENCES

1. Deshwal, V.K. 2021. Role of microbial diversity in ecosystem sustainability. *In* Biological Diversity (ed), Current Status and Conservation policies. Agro Environ Media, Publication Cell of Agriculture and Environmental Science Academy, INDIA, 1:242-255.

2. Osburn ED, Gaowen Y, Rilling M, Strickland M. 2023. Evaluating the role of bacterial diversity in supporting soil ecosystem functions under anthropogenic stress. ISME Commun 1–10.

3. Petters S, Söllinger A, Bengtsson M, Urich T. 2021. The soil microbial food’s web revisited with metatranscriptomics - predatory Myxobacteria as keystone taxon? ISME J 15:2665–2675.

4. Richenback H. 1999. The ecology of Myxobacteria. Environ Microbiol 1:15–21.

5. Wang C, Lv Y, Zhou L, Zhang Y, Yao Q, Zhu H. 2022. Comparative genomics of *Myxococcus* and *Pyxidicoccus*, including the description of four novel species: *Myxococcus guangdongensis* sp. nov., *Myxococcus qinghaiensis* sp. nov., Myxococcus dinghuensis sp. nov., and Pyxidicoccus xibeiensis sp. nov. Front Microbiol 13:995049.

6. Phillips KE, Shukria A, Cole SD. 2022. Concepts and conjectures concerning predatory performance of myxobacteria. Front Microbiol 13:1031346.

7. Waite DW, Chuvochina M, Pelikan C, Parks DH, Yilmaz P, Wagner M, Loy A, Naganuma T, Nakai R, Whitman WB, Hahn MW, Kuever J, Hugenholtz P. 2020. Proposal to reclassify the proteobacterial classes *Deltaproteobacteria* and *Oligoflexia*, and the phylum *Thermodesulfobacteria* into four phyla reflecting major functional capabilities. Int J Syst Evol Microbiol 70:5972–6016.

8. Oren A, Garrity GM. 2021. Valid publication of the names of forty-two phyla of prokaryotes. Int J Syst Evol Microbiol 71:005056.

9. Chambers J, Sparks N, Sydney N, Livingstone PG, Cookson AR, Whitworth DE. 2020. Comparative Genomics and Pan-Genomics of the Myxococcaceae, including a Description of Five Novel Species: *Myxococcus* eversor sp. nov., *Myxococcus llanfairpwllgwyngyllgogerychwyrndrobwllllantysiliogogogochensis* sp. nov., *Myxococcus* vastator sp. nov., Pyxidicoccus caerfyrddinensis sp. nov., and Pyxidicoccus trucidator sp. nov. Genome Biol Evol 12:2289–2302.

10. Sajedi H, Mohammadipanah F, Pashaei A. 2019. Automated identification of Myxobacterial genera using Convolutional Neural Network. Sci Rep 9:18238.

11. Keane R, Berleman J. 2016. The predatory life cycle of *Myxococcus xanthus*. Microbiol 62:1–11.

12. Muñoz-Dorado J, Marcos-Torres FJ, García-Bravo E, Moraleda-Muñoz A, Pérez J. 2016. Myxobacteria: Moving, Killing, Feeding, and Surviving Together. Front Microbiol 7:781.

13. Seef S, Herrou J, de Boissier P, My L., Brasseur G, Robert D, Jain R, Mercier R, Cascales E, Habermann B.H, Mignot T. 2021. A Tad-like apparatus is required for contact-dependent prey killing in predatory social bacteria. Elife 10:e72409–e72428.

14. Thiery S, Turowski P, Berleman JE, Kaimer C. 2022. The predatory soil bacterium *Myxococcus xanthus* combines a Tad- and an atypical type 3-like protein secretion system to kill bacterial cells. Cell Rep 40:111340.

15. Pérez J, Contreras-Moreno FJ, Muñoz-Dorado J, Moraleda-Muñoz A. 2022. Development versus predation: Transcriptomic changes during the lifecycle of *Myxococcus xanthus*. Front Microbiol 13:1004476.

16. Contreras-Moreno FJ, Pérez J, Muñoz-Dorado J, Moraleda-Muñoz A, Marcos-Torres FJ. 2024. *Myxococcus xanthus* predation: an updated overview. Front Microbiol 15:1339696.

17. Herrou J, Murat D, Mignot T. 2024. Gear up! An overview of the molecular equipment used by *Myxococcus* to move, kill, and divide in prey colonies. Curr Opin Microbiol 80:102492.

18. Herrou J, My L, Monteil CL, Bergot M, Jain R, Martinez E, Mignot T. 2025. Tad pili with adaptable tips mediate contact-dependent killing during bacterial predation. Nat Commun 16:4425.

19. Dong H, Xu X, Gao R, Li Y, Li A, Yao Q, Zhu H. 2022. *Myxococcus xanthus* R31 Suppresses Tomato Bacterial Wilt by Inhibiting the Pathogen *Ralstonia solanacearum* With Secreted Proteins. Front Microbiol 12:801091.

20. Ye X, Li Z, Luo X, Wang W, Li Y, Li R, Zhang B, Qiao Y, Zhou J, Fan J, Wang H, Huang Y, Cao H, Cui Z, Zhang R. 2020. A predatory myxobacterium controls cucumber *Fusarium* wilt by regulating the soil microbial community. Microbiome 8:49.

21. Yang Y, Tao H, Ma W, Wang N, Chen X, Wang W. 2023. Lysis profile and preference of *Myxococcus* sp. PT13 for typical soil bacteria. Front Microbiol 14:1211756.

22. Lewis WH, Tahon G, Geesink P, Sousa DZ, Ettema TJG. 2021. Innovations to culturing the uncultured microbial majority. Nat Rev Microbiol 19:225–240.

23. Wu K, Ma S, Li J, Sousa DZ, Cheng L. 2025. Culturing the uncultured anaerobes from the perspective of microbial growth curves. 2025. Trends Microbiol 34:89-97.

24. Thompson LR, Sanders JG, McDonald D, Amir A, Ladau J, Locey KJ, Prill RJ, Tripathi A, Gibbons SM, Ackermann G, Navas-Molina JA, Jansson JK, Gilbert JA, Knight R. 2017. The Earth Microbiome Project Consortium. A communal catalogue reveals Earth’s multiscale microbial diversity. Nature 551:457–463.

25. Ma B, Wang Y, Ye S., Liu S, Stirling E, Gilbert JA, Faust K, Knight R, Jansson JK, Cardona C, Röttjers L, Xu J. 2020. Earth microbial co-occurrence network reveals interconnection pattern across microbiomes. Microbiome 8:82.

26. Wang J, Wang J, Wu S, Zhang Z, Li Y. 2021. Global Geographic Diversity and Distribution of the Myxobacteria. Microbiol Spectr 9:e0001221.

27. Gloor GB, Macklaim JM, Pawlowsky-Glahn V, Egozcue JJ. 2017. Microbiome Datasets Are Compositional: And This Is Not Optional. Front Microbiol 8:2224.

28. Zhang L, Lueders T. 2017. Micropredator niche differentiation between bulk soil and rhizosphere of an agricultural soil depends on bacterial prey. FEMS Microbiol Ecol 93: fix103.

29. Zhou XW, Li SG, Li W, Jiang DM, Han K, Wu ZH, Li YZ. 2014. Myxobacterial community is a predominant and highly diverse bacterial group in soil niches. Environ Microbiol Rep 6:45–56.

30. Dai W, Wang N, Wang W, Ye X, Cui Z, Wang J, Yao D, Dong Y, Wang H. 2021. Community Profile and Drivers of Predatory Myxobacteria under Different Compost Manures. Microorganisms 9:2193.

31. Ding C, Hu W, Qi X, Hou S, Li Y, Xiong J, Chen T, Zhang X. 2024. Revealing the characteristics of myxobacterial communities in rhizosphere and non-rhizosphere soils of halophytic plants following wetland degradation using high-throughput sequencing. Rhizosphere 32:100972.

32. Faust K, Raes J. 2012. Microbial interactions: from networks to models. Nat Rev Microbiol 10:538–50.

33. Peschel S, Müller CL, von Mutius E, Boulesteix AL, Depner M. 2021. NetCoMi: network construction and comparison for microbiome data in R. Brief Bioinform 22:bbaa290.

34. Kodera SM, Das P, Gilbert JA, Lutz HL. 2022. Conceptual strategies for characterizing interactions in microbial communities. iScience 25:103775.

35. Suzuki SS, Baba YG, Toju H. 2023. Dynamics of species-rich predator-prey networks and seasonal alternations of core species. Nature Ecol Evol 2023 7:1432–1443.

36. Brockhurst M, Koskella B. 2013. Experimental coevolution of species interactions. Trends Ecol Evol 367-375.

37. Nair RR, Vasse M, Wielgoss S, Sun L, Yu YN, Velicer GJ. 2019. Bacterial predator-prey coevolution accelerates genome evolution and selects on virulence-associated prey defences. Nat Commun 10:4301.

38. Dai W, Liu Y, Yao D, Wang N, Shao J, Ye X, Cui Z, Zong H, Tian L, Chen X, Wang H. 2024. Biogeographic distribution, assembly processes and potential nutrient cycling functions of myxobacteria communities in typical agricultural soils in China. Sci Total Environ 906:167255.

39. McDonald D, Jiang Y, Balaban M, Cantrell K, Zhu Q, Gonzalez A, Morton JT, Nicolaou G, Parks DH, Karst SM, Albertsen M, Hugenholtz P, DeSantis T, Song SJ, Bartko A, Havulinna AS, Jousilahti P, Cheng S, Inouye M, Niiranen T, Jain M, Salomaa V, Lahti L, Mirarab S, Knight R. 2024. Greengenes2 unifies microbial data in a single reference tree. Nat Biotechnol 42:715–718.

40. Bolyen E, Rideout JR, Dillon MR, Bokulich NA, Abnet CC, Al-Ghalith GA, Alexander H, Alm EJ, Arumugam M, Asnicar F, Bai Y, Bisanz JE, Bittinger K, Brejnrod A, Brislawn CJ, Brown CT, Callahan BJ, Caraballo-Rodríguez AM, Chase J, Cope EK, Da Silva R, Diener C, Dorrestein PC, Douglas GM, Durall DM, Duvallet C, Edwardson CF, Ernst M, Estaki M, Fouquier J, Gauglitz JM, Gibbons SM, Gibson DL, Gonzalez A, Gorlick K, Guo J, Hillmann B, Holmes S, Holste H, Huttenhower C, Huttley GA, Janssen S, Jarmusch AK, Jiang L, Kaehler BD, Kang KB, Keefe CR, Keim P, Kelley ST, Knights D, Koester I, Kosciolek T, Kreps J, Langille MGI, Lee J, Ley R, Liu YX, Loft E, Lozupone C, Maher M, Marotz C, Martin BD, McDonald D, McIver LJ, Melnik AV, Metcalf JL, Morgan SC, Morton JT, Naimey AT, Navas-Molina JA, Nothias LF, Orchanian SB, Pearson T, Peoples SL, Petras D, Preuss ML, Pruesse E, Rasmussen LB, Rivers A, Robeson MS 2nd, Rosenthal P, Segata N, Shaffer M, Shiffer A, Sinha R, Song SJ, Spear JR, Swafford AD, Thompson LR, Torres PJ, Trinh P, Tripathi A, Turnbaugh PJ, Ul-Hasan S, van der Hooft JJJ, Vargas F, Vázquez-Baeza Y, Vogtmann E, von Hippel M, Walters W, Wan Y, Wang M, Warren J, Weber KC, Williamson CHD, Willis AD, Xu ZZ, Zaneveld JR, Zhang Y, Zhu Q, Knight R, Caporaso JG. 2019. Reproducible, interactive, scalable and extensible microbiome data science using QIIME 2. Nat Biotechnol 37:852–857.

41. Estaki M, Jiang L, Bokulich NA, McDonald D, González A, Kosciolek T, Martino C, Zhu Q, Birmingham A, Vázquez-Baeza Y, Dillon MR, Bolyen E, Caporaso JG, Knight R. QIIME 2 Enables 2020. Comprehensive End-to-End Analysis of Diverse Microbiome Data and Comparative Studies with Publicly Available Data. Curr Protoc Bioinformatics 70:e100.

42. McMurdie PJ, Holmes S. 2013. phyloseq: An R Package for Reproducible Interactive Analysis and Graphics of Microbiome Census Data. PLOS ONE 8: e61217.

43. Chen X, He Z, Zhao J, Liao M, Xue Y, Zhou J, Hoare R, Monaghan SJ, Wang N, Pang H, Sun C. 2022. Metagenomic Analysis of Bacterial Communities and Antibiotic Resistance Genes in Penaeus monodon Biofloc-Based Aquaculture Environments. Front Mar Sci 8:2296–7745.

44. Segata N, Izard J, Waldron L, Gevers D, Miropolsky L, Garrett WS, Huttenhower C. 2011. Meta-genomic biomarker discovery and explanation. Genome Biol 12:R60.

45. Cao Y, Dong Q, Wang D, Zhang P, Liu Y, Niu C. 2022. microbiomeMarker: an R/Bioconductor package for microbiome marker identification and visualization. Bioinformatics 38:4027–4029.

46. Barberán A, Bates ST, Casamayor EO, Fierer N. 2012. Using network analysis to explore co-occurrence patterns in soil microbial communities. ISME J 6:343–51.

47. Williams RJ, Howe A, Hofmockel KS. 2014. Demonstrating microbial co-occurrence pattern analyses within and between ecosystems. Front Microbiol 5:358.

48. Wu-Chuang A, Hartmann D, Maitre A, Mateos-Hernández L, Frantová H, Urbanová V, Obregon D, Cabezas-Cruz A, Perner J. 2023. Variation of bacterial community assembly over developmental stages and midgut of *Dermanyssus gallinae*. Microb Ecol 86 :2400–2413

49. Maitre A, Kratou M, Corona-Guerrero I, Abuin-Denis L, Mateos-Hernández L, Mosqueda J, Almazan C, Said MB, Piloto-Sardiñas E, Obregon D, Cabezas-Cruz A. 2024. Differential interactions of *Rickettsia* species with tick microbiota in *Rh. sanguineus* and *Rh. turanicus*. Sci Rep 14:20674.

50. Aitchison J. 1982. The statistical analysis of compositional data. J.R Stat Soc B Methodol 44:139–60.

51. Shannon P, Markiel A, Ozier O, Baliga NS, Wang JT, Ramage D, Amin N, Schwikowski B, Ideker T. 2003. Cytoscape: a software environment for integrated models of biomolecular interaction networks. Genome Res 13:2498–504.

52. Ruhnau B. 2000. Eigenevector-centrality—a node-centrality? Soc Networks 22:357–365.

