## Supplemental figures for "Evidence for a central ecological function of predatory Myxococcota in soil ecosystems"

**Figure S1.** Venn diagrams comparing the proportions of microbial (A) and Myxococcota (B) taxa between field soil and sand biofilter environments.

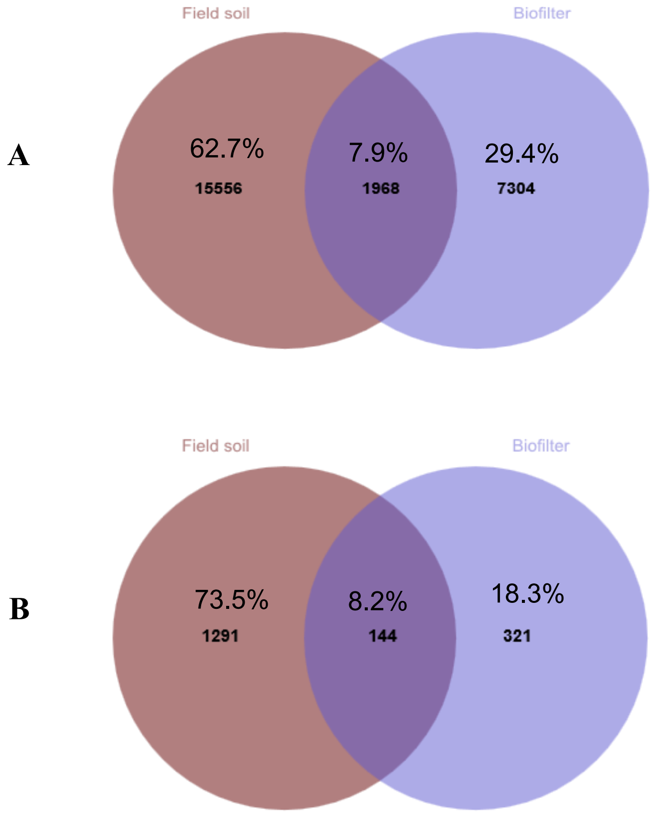

**Figure S2.** Patterns of microbial phyla identified across modules in field soil network as measured by proportions of taxa and of degree and eigen centralities.

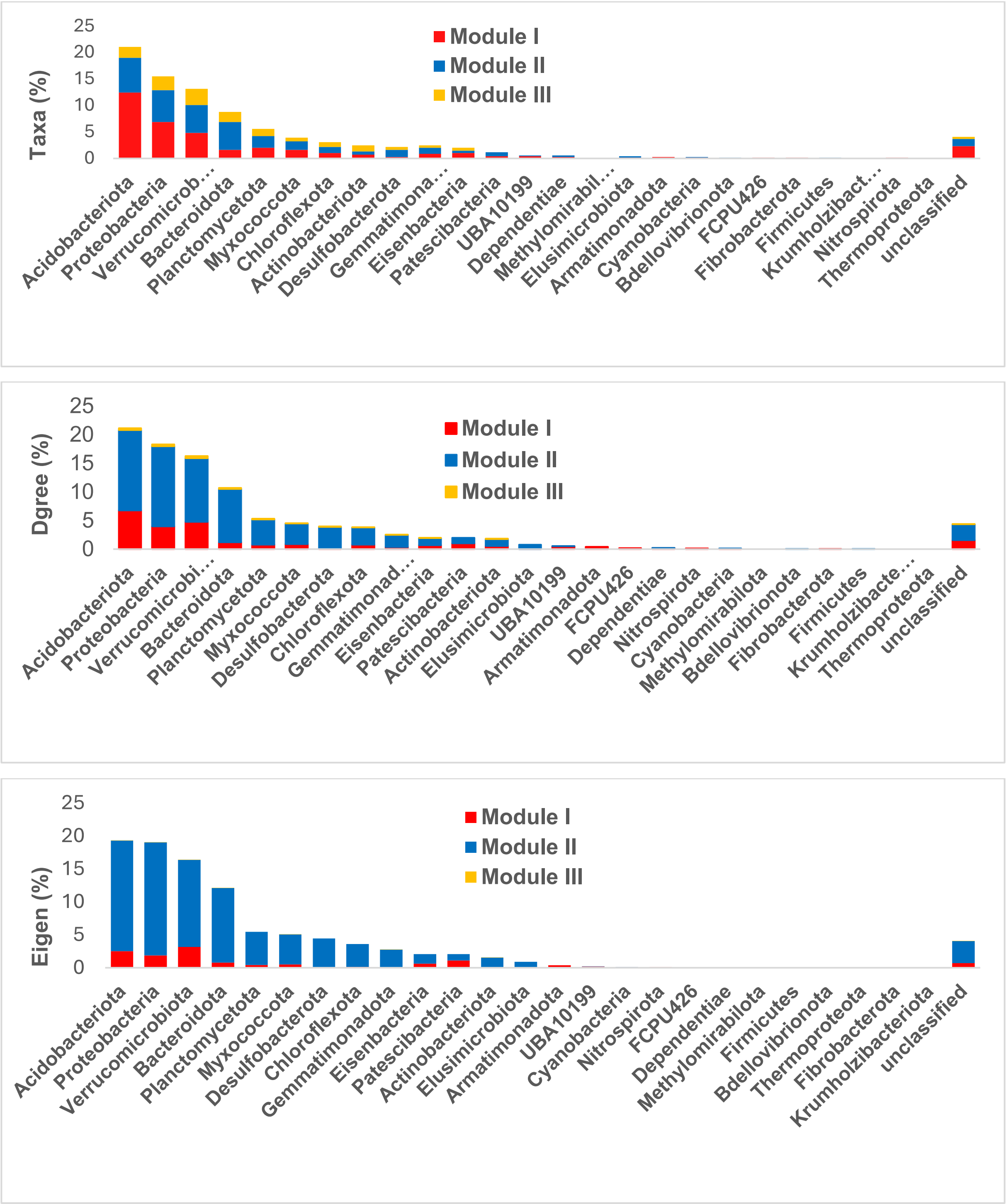

**Figure S3.** Distribution of neighboring phyla of Myxococcota by modules in field soil subnetwork as measured by degree centrality.

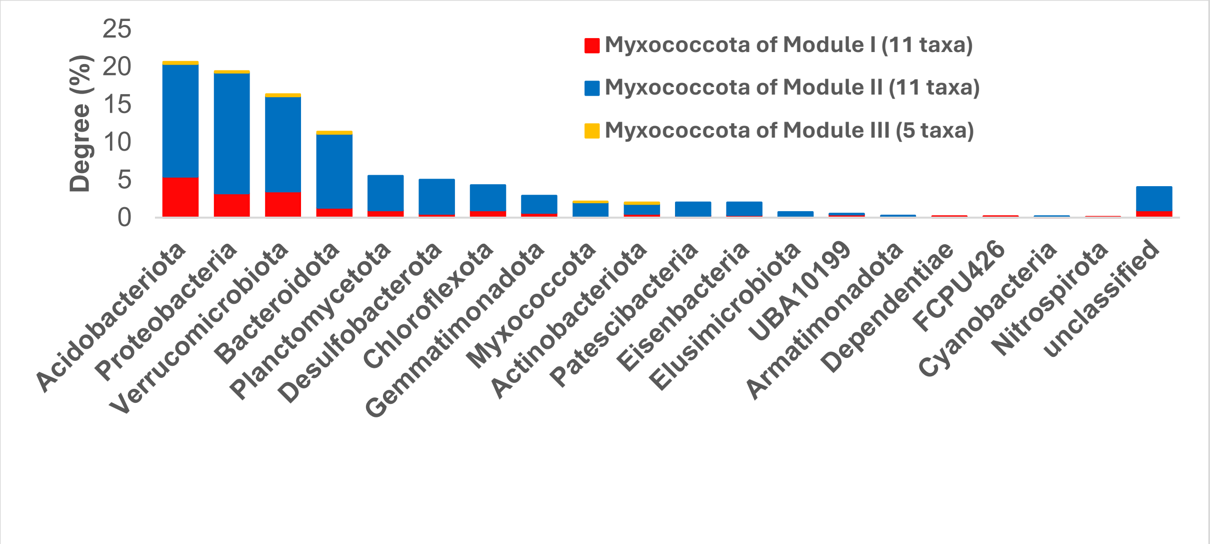

**Figure S4.** Patterns of microbial phyla detected across modules in sand biofilter network as measured by proportions of taxa and of degree and eigen centralities.

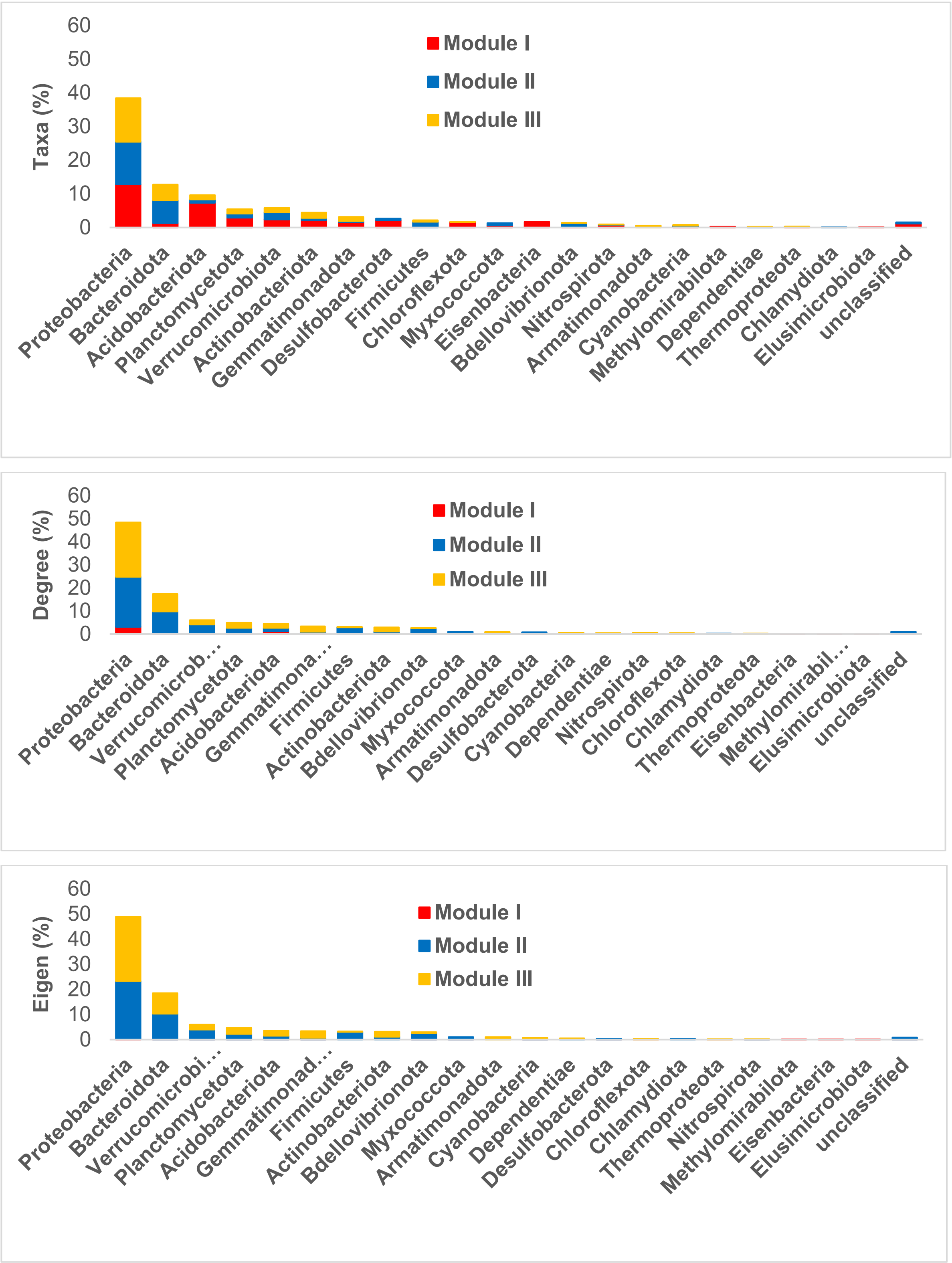

**Figure S5.** Distribution of neighboring phyla of Myxococcota by modules in sand biofilter subnetwork as measured by degree centrality.

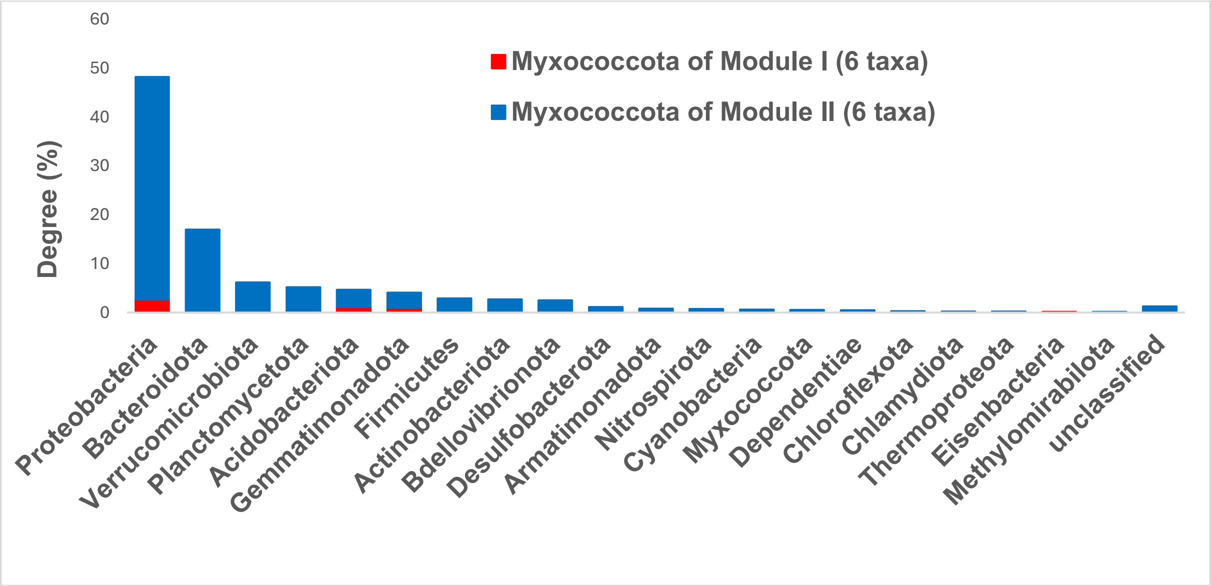
